# Decoupling Lineage and Intrinsic Information in Single-Cell Lineage Tracing Data with Deep Disentangled Representation Learning

**DOI:** 10.64898/2026.03.10.710716

**Authors:** Yuhong Wen, Jiankang Xiong, Fuzhou Gong, Liang Ma, Lin Wan

## Abstract

Single-cell RNA sequencing combined with lineage tracing technologies provides rich opportunities to study development and tumor evolution, yet existing computational methods struggle to disentangle intrinsic transcriptional states from lineage-driven effects. We introduce DeepTracing, a deep generative framework that integrates disentangled representation learning with lineage-aware Gaussian processes to explicitly separate intrinsic cellular variation from lineage constraints. The model constructs a layered latent space and enforces independence via Total Correlation regularization, producing intrinsic, lineage, and unified embeddings. Across extensive benchmarks, DeepTracing consistently outperforms existing approaches. In TedSim simulations, it achieves superior clustering of cell states and effectively recovers phylogenetic structure, surpassing original expression and scVI. Applied to mouse tumor lineage-tracing data, DeepTracing attains higher ARI/NMI for tumor-type classification than scVI and PORCELAN, accurately separating primary and metastatic tumors and recovering known trajectories such as early lymph-node divergence and liver-to-kidney cross-seeding. In larger datasets, it maintains strong performance while preserving both transcriptomic continuity and lineage fidelity. DeepTracing also reconstructs continuous developmental trajectories in mouse ventral midbrain, isolating temporal effects from intrinsic differentiation. These results establish DeepTracing as a scalable and interpretable framework for analyzing multimodal single-cell data in tumor progression.

**Code availability:** The source code is publicly available at https://github.com/Yuhong-Wen/DeepTracing.

## 1 Introduction

Recent advances in single-cell RNA sequencing (scRNA-seq) combined with lineage tracing technologies have enabled the simultaneous capture of cellular gene expression profiles and lineage information [1,2,3]. These multimodal datasets provide unprecedented opportunities to study developmental processes and tumor progression in complex multicellular systems [4], revealing not only snapshots of transient cellular states but also their ancestral relationships and dynamic evolution. Various computational methods have been developed to analyze such data. Existing approaches primarily focus on trajectory inference [5,6,7,8,9], lineage tree reconstruction [10,11,12], or the recovery of ancestral expression states [13], significantly advancing our ability to explore developmental processes from multimodal single-cell measurements.

The transcriptional characteristics of cells are simultaneously shaped by both intrinsic properties and extrinsic lineage relationships. Intrinsic properties refer to the type-specific characteristics of cells and their cell states [14], while extrinsic lineage relationships reflect the genetic relationships and division of labor trajectories of cells throughout their developmental history, often traced through phylogenetic barcodes or epigenetic memories [15]. These two factors interweave, jointly determining the current transcriptional state and functional characteristics of cells. Decoupling their contributions to gene expression is crucial for understanding the mechanisms of cell fate determination, the basis for maintaining identity stability, and the underlying patterns of intrinsic disorder in related diseases.

Decoupling the contributions of these two factors presents fundamental challenges. Cells that share the same lineage often cluster together both spatially and molecularly during early development, meaning that observed gene expression patterns largely reflect their common ancestry rather than cell-autonomous programs. Moreover, even when descending from the same progenitor, different cell types or states may interpret and respond differently to lineage-derived signals, leading to heterogeneous fate decisions. Notably, most existing computational methods do not adequately address the challenge of decoupling intrinsic and lineage-driven variation in single-cell data. Standard visualization approaches such as UMAP [16] and t-SNE [17] tend to organize cells based on their current transcriptional states, which can reveal expression continuity but often obscures the underlying lineage tree structure. There is always a trade-off: emphasizing expression continuity maintains smooth gradients of cellular states but blurs or erases the true lineage structure [18,19], whereas preserving lineage fidelity recovers ancestral connections but disrupts continuous expression-defined trajectories. Efforts to integrate both aspects often lead to entangled representations of lineage and state, yielding results that are difficult to interpret and potentially misleading for downstream analyses [20,21].

The challenge of this task is further compounded by the biological complexity of lineage-state relationships. In some subtrees, lineage and transcriptional states are tightly coupled, resulting in descendant cells with homogeneous expression profiles [15]; in others, expression dynamics become decoupled from lineage, as intrinsic transcriptional programs drive divergent fate outcomes despite shared ancestry [22]. Between these extremes, intermediate coupling produces mixed phenotypes, reflecting competition between lineage constraints and cell-autonomous adaptation.

To overcome these limitations, we propose DeepTracing, a deep learning-based multimodal model that leverages disentangled representation learning to explicitly decouple and integrate lineage and state information. DeepTracing constructs a layered latent space by combining standard Gaussian priors to capture intrinsic cellular properties and Gaussian process priors to model lineage constraints, with the Total Correlation penalty [23] that explicitly enforces statistical independence between the two latent portions, thus ensuring their decoupling. This design produces three complementary embeddings: an intrinsic embedding that reflects gradual changes in cell identity and state, a lineage embedding that preserves hierarchical structure, and a unified embedding that integrates both dimensions into a coherent latent representation. In this way, DeepTracing maintains continuity of expression-defined states while simultaneously recovering lineage fidelity, effectively resolving the long-standing trade-off in existing methods.

We validated the disentangled modeling capability of DeepTracing using synthetic data generated by TedSim [24], where the model simultaneously recovered ground-truth cell states and lineage subtrees from leaf-node observations. We then applied DeepTracing to CRISPR-based lineage-traced lung tumor datasets, in which the unified embedding cleanly separated primary and metastatic lesions, recapitulated experimentally validated metastatic routes, and enabled the discovery of metastasis-associated gene programs. Finally, in a mouse ventral midbrain dataset spanning multiple embryonic time points, DeepTracing disentangled temporal effects from intrinsic cellular states, reconstructing a smooth continuum of neuronal differentiation while preserving sampling-time information in a dedicated latent layer. Together, these results demonstrate that DeepTracing provides a flexible and scalable framework for analyzing high-dimensional multimodal single-cell data, yielding representations that are both biologically interpretable and well suited for downstream tasks such as clustering, trajectory inference and marker gene discovery.

## 2 Results

### 2.1 DeepTracing

We introduce DeepTracing, a deep generative model that leverages a disentangled representation learning framework to effectively integrate and decouple scRNA-seq and lineage tracing data. Its core objective is to utilize a Variational Auto-Encoder (VAE) to learn a structured latent representation, enabling the effective separation of intrinsic cell state variation and lineage-driven effects.

The overall architecture of DeepTracing is summarized in Fig.1A. The model accepts two primary inputs: the single-cell gene expression matrix and the corresponding lineage information (e.g., cell barcoding data or a lineage tree structure). These inputs are mapped by an encoder to infer a set of latent variables. The key feature of the model is the disentanglement mechanism applied to the latent variables, which yields two statistically independent latent factors: the intrinsic variables and the lineage variables. These two decoupled factors are subsequently passed to a decoder for the reconstruction of the original gene expression matrix, which drives the reconstruction losses.

**Figure 1.**
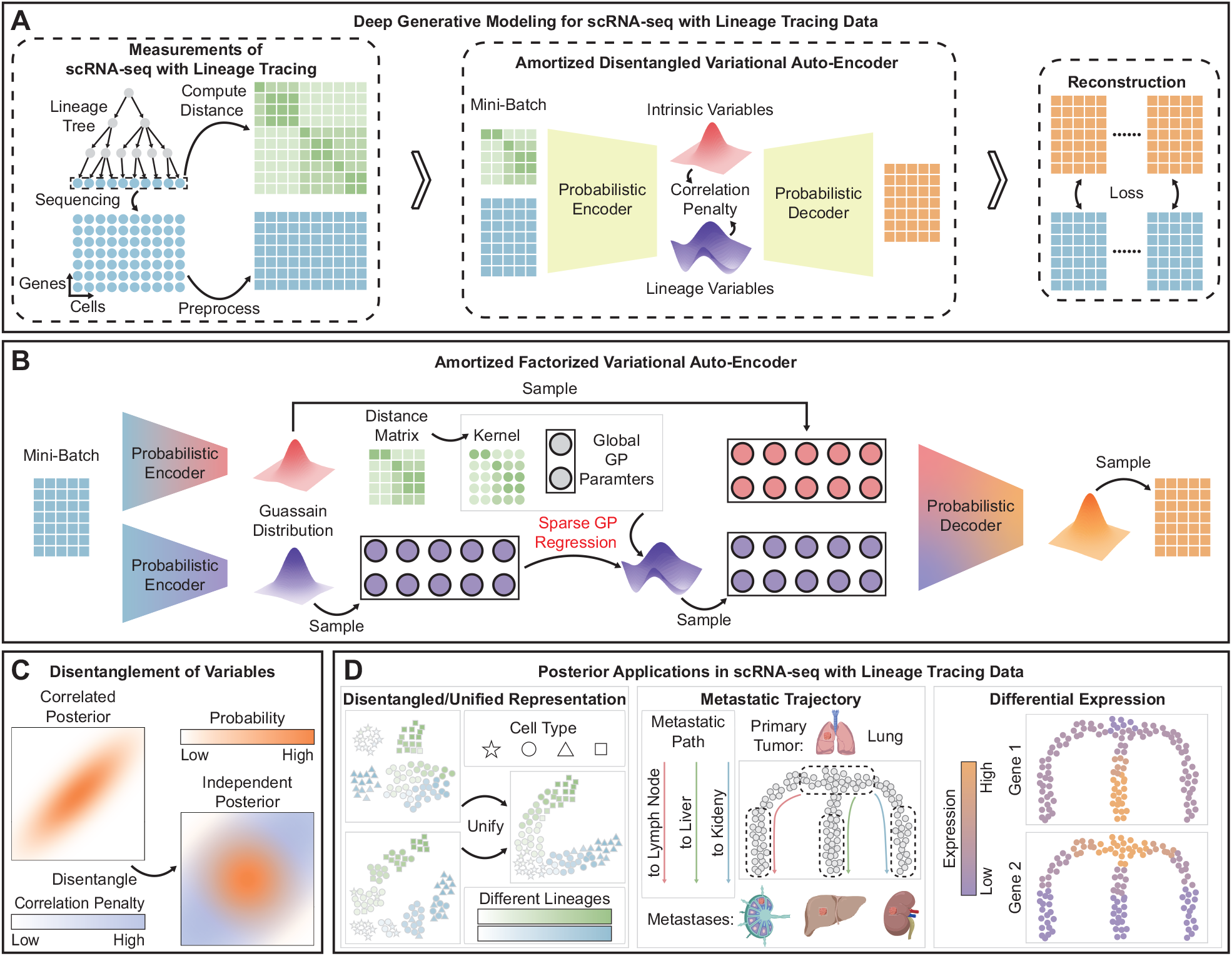
Overview of the DeepTracing model. (A) The overall architecture of DeepTracing, which illustrates the computational flow from the input of scRNA-seq and lineage tracing data, through the encoder, the layered latent space, and reconstruction via the decoder. (B) The structural design of the DeepTracing latent space, showing the factorization of the latent space into the intrinsic state embedding and the lineage embedding, along with their distinct prior constraints. (C) A schematic representation of the decoupling mechanism, comparing the changes in correlation of the posterior distributions of the latent variables before and after applying the TC penalty constraint. (D) The disentangled/unified representations generated by the model and their downstream biological applications. (The schematics for the organs in the Metastatic Trajectory panel are sourced from BioGDP.com [25].)

The latent space is explicitly designed as a layered structure (Fig.1B) to enforce the separation of biological variation. The total latent space, is factorized into two components: the intrinsic state embedding, which is responsible for capturing localized cellular properties like cell type and cell identity; and the lineage embedding, which captures the global constraints imposed by ancestral relationships and developmental history. The priors for these components are distinct: the intrinsic state embedding is governed by a standard Gaussian prior, while the lineage embedding is specifically constrained by a Gaussian Process (GP) prior, where the kernel matrix is derived from the calculated lineage distance between cells. Furthermore, to deal with large-scale datasets, DeepTracing employs a scalable approach that combines amortized variational inference with sparse Gaussian process regression using inducing points.

The mechanism to enforce statistical independence between these two factors is crucial, as illustrated in Fig.1C. Without explicit constraints, the posterior distributions of the two latent variables often remain correlated. DeepTracing overcomes this by applying a Total Correlation (TC) penalty to explicitly minimize the mutual information between the latent factors, leading to a statistically independent latent space, which is essential for accurate biological interpretation.

The utility of these disentangled representations for downstream biological analysis is summarized in Fig.1D. The model provides three distinct representations of the data: the lineage embedding, the intrinsic embedding, and the unified embedding. The lineage embedding recovers the true ancestral structure and enhances lineage fidelity. The intrinsic embedding distinguishes distinct cellular states. The resulting unified embedding, which integrates both decoupled factors, provides a comprehensive basis for further analyses, including the inference of metastatic trajectories and conducting differential expression analysis to identify genes associated with either state transition or lineage progression.

### 2.2 DeepTracing effectively decouples variation

We benchmark DeepTracing with TedSim [24] simulation dataset (Supplementary Note 1). The TedSim model simulates cell state transition process through cell division events, and coupled with a barcode generation process. The simulated cell lineages are represented as a binary tree capturing division history, with cell states assigned according to a predefined cell state tree that specifies permissible transitions toward terminal states [24]. Since experimental observations typically capture only terminal cells (leaf nodes) of the lineage tree, we focused our analysis on the profiles of leaf nodes, and applied the scaled Hamming distance defined in [6] to barcodes. The lineage tree structure can be extracted from the simulation as ground truth. We partitioned this tree into subtrees rooted at nodes of depth 4. The terminal cells were assigned to corresponding subtree as a representation of true lineage identity (Fig.2A).

**Figure 2.**
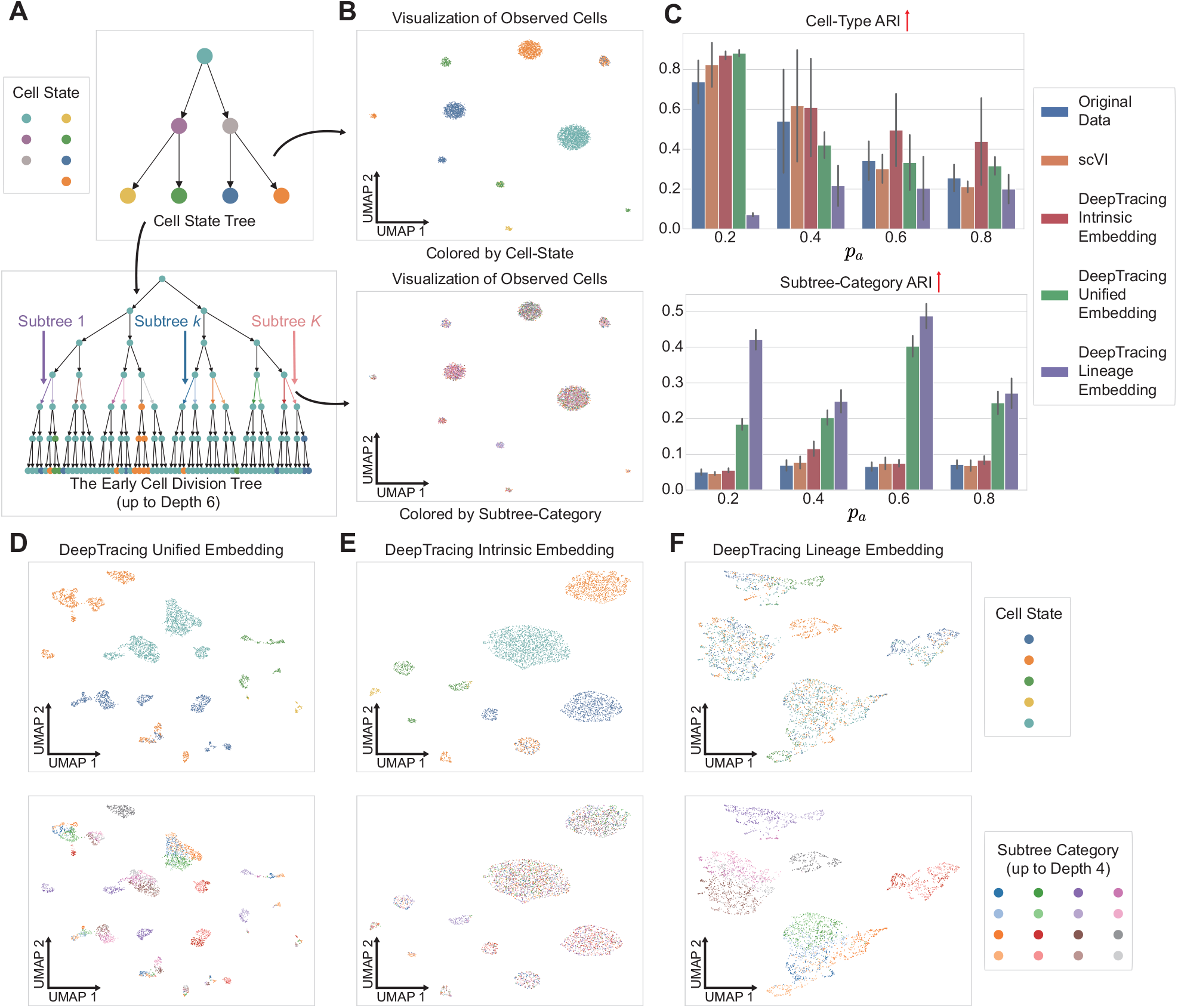
DeepTracing performance on the TedSim simulated dataset. (A) The underlying cell state tree (top) and the simulated early cell division tree (bottom, up to depth 6) generated by TedSim. The cell states and subtree categories are indicated by color. (B) UMAP visualization of the original simulated gene expression data when the simulation parameter *p*_*a*_ = 0.2, colored by cell state (top) and subtree category (bottom), respectively. (C) Celltype ARI (top) and subtree-category ARI (bottom) metrics for the learned embeddings across various *p*_*a*_ values. (D-F) UMAP visualizations of the learned embeddings from DeepTracing when *p*_*a*_ = 0.2: (D) unified embedding, (E) intrinsic embedding, and (F) lineage embedding. The upper row in (D-F) is colored by cell state, and the lower row is colored by subtree category (up to depth 4).

The UMAP visualization shows that in the original gene expression space, cells primarily form distinct clusters based on their cell-states, while their lineage identities (subtrees) were mixed within each cluster (Fig.2B), revealing the dominance of state information at the transcriptome level. We applied DeepTracing to this simulation data, which resulted in three latent representations. We then performed *K* -means[29] clustering on each of representation as well as on the original expression profile and the latent space derived from scVI [28]. We adopted the Adjusted Rand Index (ARI) [30] and Normalized Mutual Information (NMI) [31] (Supplementary Note 2) to evaluated the clustering results against ground-truth cell state labels and lineage subtree categories. DeepTracing successfully disentangled cell-state variability and lineage identity (Fig.2C-F; Supplementary Fig.S1A): the intrinsic embedding captured cell state features with highest ARI and NMI scores than those obtained from the original gene expression space or the scVI embedding. The lineage embedding showed the ability of separating lineage subtree, with significantly increased ARI and NMI as compared to others. The unified embedding, aggregate both types of information, which showed good separation of both cell-state and lineage identity. Moreover, DeepTracing also demonstrated superior performance across multiple unsupervised clustering metrics (Supplementary Note 2), further validating its effectiveness (Supplementary Fig.S1B).

To validate the robustness of DeepTracing, we conducted experiments on a key parameter, *p*_*a*_, in the TedSim simulation. *p*_*a*_ represents the probability of cell asymmetric division. A higher *p*_*a*_ value leads to more frequent state transitions in the simulated lineage tree [24]. We set *p*_*a*_ values to 0.2, 0.4, 0.6, and 0.8, simulating four datasets, and performed ten replicate experiments for each method. The results show that as *p*_*a*_ increases, and thus the complexity of the generated data rises, DeepTracing is slightly affected but remains generally stable overall (Fig.2C; Supplementary Fig.S1).

It is worth noting that methods for analyzing lineage data, such as PORCELAN [26], which uses a triplet loss through the variational autoencoder [27] to strongly aggregate cells from the same lineage subtree, often require constructing full distance structures and triplets among cells, thereby making it computationally expensive and time-consuming. The simulated datasets, each consisting of nearly 4,000 cells, present a challenge for PORCELAN, as it struggles to produce valid results under these conditions. In contrast, while DeepTracing also incorporates similar distance structures through Gaussian processes, it uses an amortized optimization approach that allows for iterative training on small batches of data, significantly improving both computational power and efficiency.

### 2.3 DeepTracing verifies potential tumor metastasis trajectories

To verify the efficacy of DeepTracing in parsing tumor heterogeneity and evolutionary relationships. We applied DeepTracing to a paired lineage tracing and scRNA-seq dataset (Supplementary Note 1) of a lung cancer mouse model [32]. This mouse model leverages a CRISPR/Cas9-based lineage tracing system [33], in which cells accumulate mutations in barcode sequences during cell division. These accumulated mutations enable the inference and reconstruction of high-resolution lineage trees of the terminal cells [34].

We first focused on a tumor-metastasis sample 3515_Lkb1_T1, which contains both primary and metastatic sites comprising 1,878 cells. We found that both our lineage and unified embeddings achieved higher ARI and NMI for tumor type classification compared to the original feature space and scVI, and outperform the PORCELAN’s representation (Fig.3A). As shown in Fig.3B, primary and metastatic tumor cells are intermixed with poor separation in the original expression UMAP space. DeepTracing showed clear segregation of primary and metastasis tumors through its unified embedding, distinct tumor clones partitioned following a gradual transition path. Specifically, the Kidney metastases K2 and K3 were mostly mixed within the same cluster, indicating they are derived from a common seeding event. By contrast, samples from tumor K1 were separated into two distant clusters, one grouped with L1, K2 and K3, while the other grouped with L2. This representation aligns with the metastasis history, where K1 experienced cross-seeding events with mixed ancestry from both distinct liver sites. In sum, the joint embedding derived from DeepTracing faithfully recapitulates both the clonal structure and the metastatic history, including early lymphatic spread and liver-to-kidney cross-seeding, aligning well with experimentally validated evolutionary routes (Fig.3C), and confirming that the latent representations learned by DeepTracing are biologically meaningful. In contrast, the latent space derived from PORCELAN fragments the same data into discrete tumor-type clusters, failing to preserve the continuous spectrum of cellular states (Supplementary Fig.S2A).

**Figure 3.**
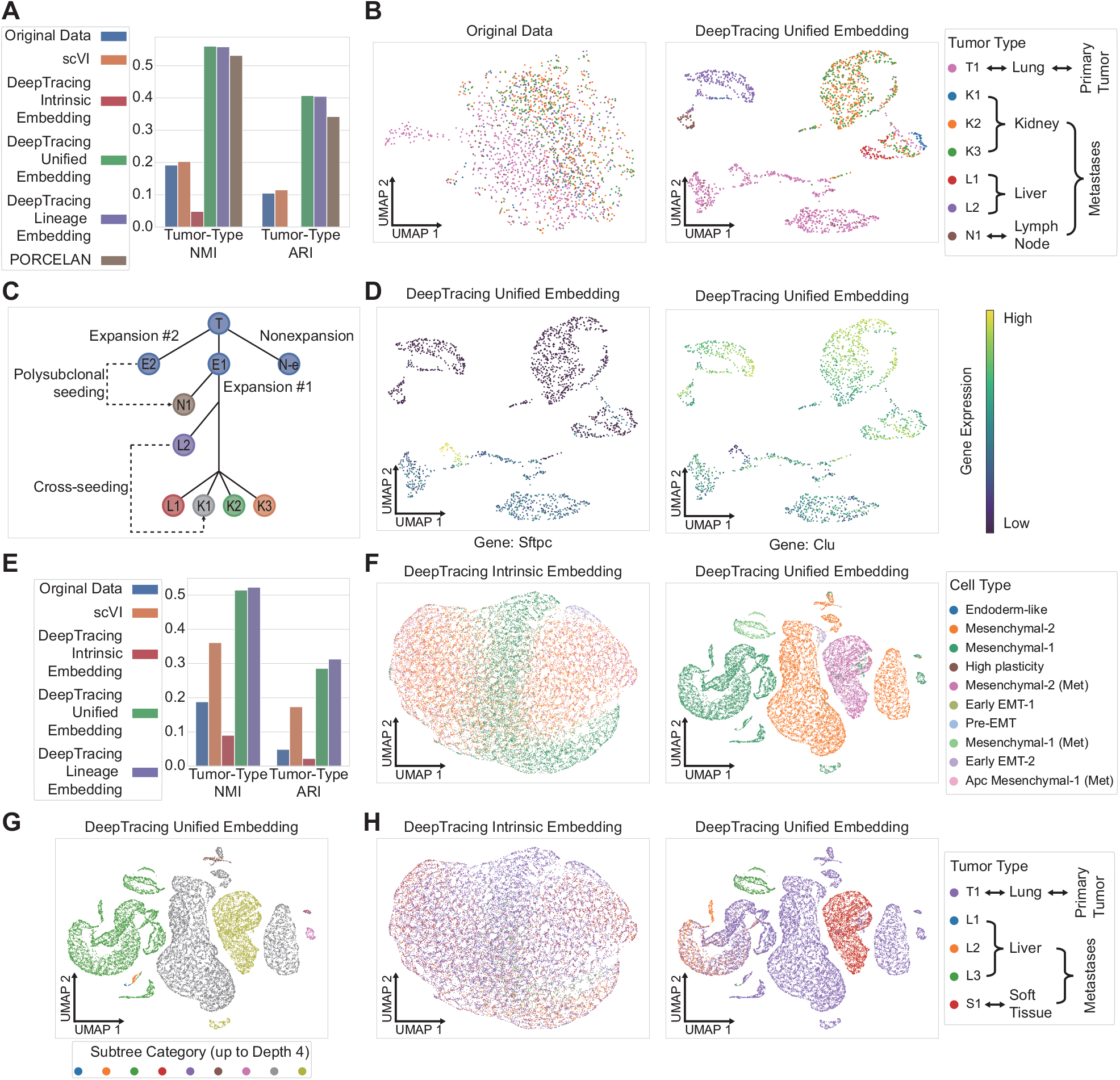
DeepTracing performance on the tumor metastasis datasets. (A-D) Results for the 3515_Lkb1_T1 family dataset: (A) Tumor-type NMI and Tumor-type ARI metrics for the learned embeddings. (B) UMAP visualization of the original gene expression data (left) and the DeepTracing unified embedding (right), colored by tumor type. (C) A model summarizing the corresponding metastatic behaviors. (D) UMAP visualization of the DeepTracing unified embedding, colored by expression levels of the top two marker genes identified by CCA: *Sftpc* (left) and *Clu* (right). (E-H) Results for the 3724_N1_T1 family dataset: (E) Tumor-type NMI and Tumor-type ARI metrics for the learned embeddings. (F) UMAP visualization of the DeepTracing intrinsic embedding (left) and the DeepTracing unified embedding (right), colored by cell type. (G) UMAP visualization of the DeepTracing unified embedding, colored by subtree category. (H) UMAP visualization of the DeepTracing intrinsic embedding (left) and the DeepTracing unified embedding (right), colored by tumor type.

In addition, DeepTracing’s embeddings can be used for genetic analysis. We applied Canonical Correlation Analysis (CCA) [35] between gene expression and the unified embedding to identify metastasis-related genes. The top two genes with the highest Canonical Correlation are *Sftpc* and *Clu. Sftpc*, a marker for the alveolar type 2 (AT2)-like cell type that represents the initial state of the tumor [26,32,36], is predominantly expressed in the primary tumor and downregulated in metastases, while *Clu* [37] is upregulated in metastases (Fig.3D), which shows that DeepTracing can identify genes associated with metastasis.

### 2.4 DeepTracing reveal tumor mestatisit in large-scale lineage dataset

We further extended our analysis to a larger tumor sample 3724_NT_T1, which contained over 20,000 cells, including the primary tumor, three liver metastases, and one soft tissue metastasis, among them, liver and soft tissue metastases originated from distinct expanded subclones occupying disparate spatial niches and distant phylogenetic positions [32]. In this context, both DeepTracing’s lineage and unified embeddings achieved higher ARI and NMI for tumor type classification than the original feature space or scVI, as they explicitly incorporate lineage relationships (Fig.3E). Notably, the intrinsic embedding grouped mesenchymal cells of the same subtype from diverse sites (including metastatic and non-metastatic samples) into shared clusters, highlighting their common molecular phenotype as a consistent cell type, on the other hand, the unified embedding, which incorporates lineage information (Fig.3F), organized cells into subtrees that correlated strongly with tumor types (Fig.3G, H). This result demonstrated DeepTracing’s ability of providing multi-faceted and complimentary representations of tumor heterogeneity. This integrated, multi-faceted view of heterogeneity was further evidenced by the distinct expression patterns of key genes: *lgfbp5* in liver-metastasized tumors and *Spp1* in primary tumors (downregulated in metastasis), as clearly visualized in the unified embedding (Supplementary Fig.S2B). Additionally, datasets of this scale present a prohibitive computational challenge for methods such as PORCELAN.

These results further validate that DeepTracing constructs a biologically informative space where both developmental lineage and functional state are coherently represented. Its ability to consistently resolve primary-metastasis separation across different samples highlights its generalizability and utility in uncovering tumor evolutionary patterns.

### 2.5 DeepTracing recovers the continuous developmental trajectory of cells from multiple time points

We also utilized a dataset (Supplementary Note 1) of ventral midbrain tissues from mouse embryos during the embryonic stage [38]. The original experiment introduced a mutation marker on day 9.5 (E9.5) of the embryo and isolated the ventral midbrain tissues on days 11.5 (E11.5) and 15.5 (E15.5), conducting single-cell transcriptome sequencing and reading barcodes. A total of approximately 50,000 cells were obtained, covering radial glial (Rgl), neuroblast (Nb), dopaminergic (DA), glutamatergic (GLU), GABAergic neurons, and various intermediate precursor subpopulations [38].

In this study, we integrated a subset of data from two time points (E11.5 and E15.5), encompassing over 8,000 cells. By incorporating a temporal decoupling strategy into our analysis, we were able to separate time-related effects from the intrinsic cellular states. In the resulting intrinsic embedding, cells from both time points form a continuous developmental continuum without clear separation by time (Fig.4A), indicating a smooth transition along the differentiation trajectory. In contrast, cells in the original feature space, despite being identical cell type, were clearly separated by their time points. Meanwhile, in the temporal related embedding, cells are strictly organized according to their sampled developmental stages, confirming that the temporal information has been effectively disentangled and represented (Fig.4B). This demonstrates that our model successfully disentangles the underlying continuous differentiation process from the temporal context, allowing the intrinsic structure of cell states to be visualized independently of sampling time.

**Figure 4.**
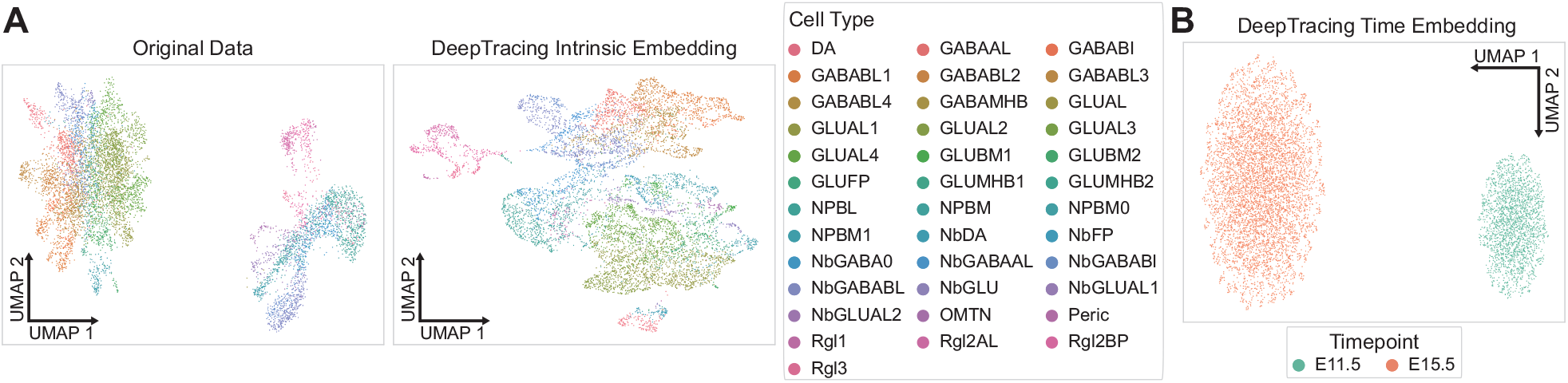
DeepTracing performance on the continuous developmental dataset. (A) UMAP visualization of the original gene expression data (left) and the DeepTracing intrinsic embedding (right), colored by cell type. (B) UMAP visualization the DeepTracing time embedding, colored by time point.

## 3 Conclusion

In this study, we have proposed DeepTracing, a powerful deep learning-based method that effectively decouples and integrates both lineage and state information from single-cell data. By leveraging Gaussian processes and disentangled representation learning, DeepTracing constructs a biologically meaningful latent space where both developmental lineage and functional state are coherently represented. Across synthetic benchmarks and multiple real datasets, DeepTracing consistently achieved superior performance in jointly modeling lineage and transcriptional variation. On TedSim simulations, its intrinsic and lineage embeddings more accurately recovered ground-truth cell states and lineage subtrees than conventional latent-variable models, while its unified embedding provided a balanced view of both sources of variation. In lung tumor lineage-tracing data, DeepTracing resolved the intermingling of primary and metastatic cells, reconstructed clonal structure and cross-seeding events that were concordant with experimentally inferred metastatic histories, and facilitated the identification of metastasis-associated marker genes. In embryonic ventral midbrain, the model decoupled sampling time from differentiation state, revealing continuous developmental trajectories that were obscured in the original expression space. These empirical results highlight the models capacity to capture complex biological structure in a wide range of experimental settings.

The application of DeepTracing to diverse datasets has shown its robustness and flexibility in capturing both broad biological structures, such as clonal relationships, and finer-grained features like gene expression variation. Its ability to maintain the continuity of cellular states while preserving lineage fidelity addresses the long-standing challenge of decoupling intrinsic and lineage-driven variation in single-cell data. Moreover, the unified embeddings generated by DeepTracing provide valuable insights into tumor heterogeneity, metastasis trajectories, and the dynamic evolution of cell states, significantly improving upon traditional methods. These findings confirm that DeepTracing can effectively decouple cell-state and lineage information, offering a powerful tool for understanding the molecular underpinnings of development, disease progression, and metastasis. Its robustness and scalability position it as a promising approach for analyzing high-dimensional, multimodal single-cell data in a wide range of biological contexts.

## 4 Methods

### 4.1 Deep generative framework of DeepTracing

We propose DeepTracing, a deep generative framework designed to model single-cell gene expression data and their corresponding cellular lineage structure information. DeepTracing is based on the VAE [27] framework, but extends it with an integrated prior mechanism that incorporates GP [39] to encode lineage structural constraints between cells. Its core objective is to integrate single-cell gene expression data and lineage information, and to distinguish the gene expression changes caused by cell types from those caused by lineage structures, ultimately producing three embeddings. It specifically incorporates a TC loss function to effectively guide the disentangled learning of latent representations [23].

Let **Y** = [**y**_1_, …, **y**_*N*_]^⊤^ ∈ ℝ^*N* ×*G*^ denote the observed gene expression matrix for *N* cells and *G* genes, where each row **y**_*i*_ ∈ ℝ^*G*^ corresponds to the expression profile of a single cell. Let **X** = [**x**_1_, …, **x**_*N*_]^⊤^ denote the auxiliary information associated with the cells, such as their lineage relationships. Our model aims to learn a latent representation **Z** = [**z**_1_, …, **z**_*N*_]^⊤^ ∈ ℝ^*N* ×*L*^ that captures the underlying biological variation.

To achieve interpretability and flexibility, we decompose the latent space into two components. Specifically, the first *P* dimensions of the latent space, denoted by **Z**^1:*P*^, capture structured variation shared across cells and guided by metadata **X**, while the remaining *L* − *P* dimensions, denoted by **Z**^*P* +1:*L*^, capture cell-specific local variation. Accordingly, we place the following priors over the latent variables:

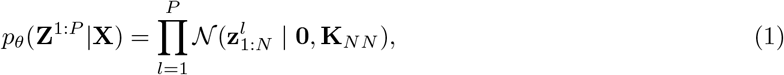

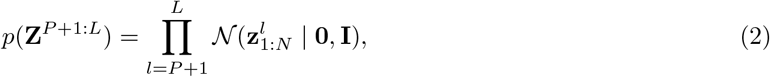

where 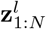 denotes the *l*th dimension across all cells and **K**_*NN*_ is a kernel matrix defined by a GP over **X**. In practice, we use lineage-aware kernels to define **K**_*NN*_ :

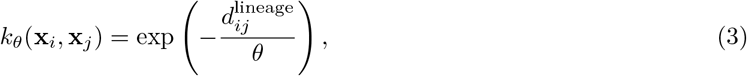

where 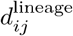 is the pairwise distance between cells *i* and *j* in lineage space, and *θ* is a learnable length-scale parameter. Two distance metrics are supported:

1. **Lineage Tree Distance**: Defined as the shortest path distance on the reconstructed lineage tree [1,34], i.e., the sum of distances from nodes *i* and *j* to their closest common ancestor: 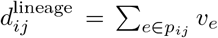, where *p*_*ij*_ is the shortest path between nodes *i* and *j* on the lineage tree, and *v*_*e*_ is the edge weight.
2. **Barcode Distance**: Defined as the Hamming distance between barcodes [6].

The inference model is parameterized by an encoder network *q*_*ψ*_(**Z** |**Y**) = 𝒩 (**w**, diag(***σ***^2^)), which outputs the mean **w** and the diagonal variance ***σ***^2^ of the approximate posterior. We apply GP-based regularization to the first *P* dimensions of the latent representation to align them with the structured prior, while the remaining dimensions are regularized toward a standard Normal prior. To explicitly promote disentanglement between lineage-induced and state-induced factors, we introduce a TC penalty. The complete training objective maximizes an augmented evidence lower bound (ELBO):

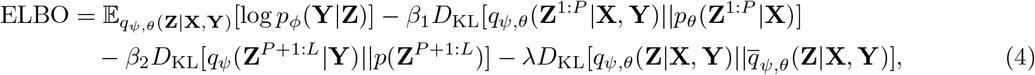

where *q*_*ψ,θ*_(**Z**|**X, Y**) is the approximate posterior, *p*_*ϕ*_(**Y**|**Z**) is the decoder network that reconstructs gene expression from the latent code, *β*_1_ and *β*_2_ are the hyperparameters that balance reconstruction accuracy with latent space regularization [40]. 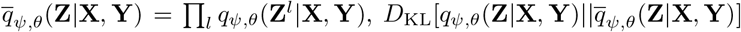 is the TC which measures dimension-wise dependence in **Z** [23]. The decoder *p*_*ϕ*_ is implemented as a probabilistic model (e.g., Negative Binomial) depending on the characteristics of the input data. During training, the model jointly learns the encoder parameters *ψ*, the decoder parameters *ϕ*, and the kernel hyperparameters *θ* in an end-to-end manner.

This integrated latent space design enables DeepTracing to effectively model both global structures such as developmental or lineage trajectories and local heterogeneity. The GP-regularized dimensions provide a smooth latent manifold that respects biological continuity, while the unstructured dimensions allow the model to capture fine-grained variability. By explicitly disentangling these two sources of variation, DeepTracing improves both interpretability and downstream utility in tasks such as trajectory inference, clustering, and marker gene identification.

### 4.2 Approximate posterior inference of DeepTracing

Training DeepTracing by maximizing the evidence lower bound (ELBO) presents significant computational challenges, mainly because GP inference requires inverting the kernel matrix **K**_*NN*_, which has a computational complexity of 𝒪 (*N* ^3^) that grows cubically with the number of samples *N*, severely limiting the model’s scalability and practical application to large datasets [41]. By leveraging amortized variational inference and sparse variational inference based on inducing points, the model’s computational complexity is reduced to 𝒪 (*bm*^2^ + *m*^3^), where *b* is the batch size and *m* is the number of inducing points (*m* ≪ *N*), significantly enhancing the model’s feasibility and computational efficiency.

The mathematical derivation focuses on the computation of the GP KL-divergence term. Consider the first *P* latent dimensions with a GP prior *p*_*θ*_(**Z**^1:*P*^ | **X**). Given the output of probabilistic encoder *q*_*ψ*_, the mean **w** and the diagonal variance ***σ***^2^, the GP-VAE combines latent representation learning with GP regression, and induces the posterior *q*_*ψ,θ*_(**Z**^1:*P*^ | **X, Y**). Based on [42], the KL loss term embedded in the GP can be written as:

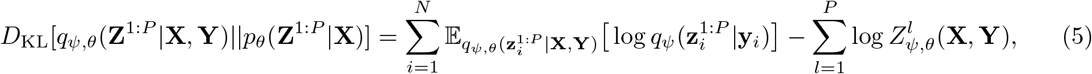

where *q*_*ψ*_(**Z**^1:*P*^ |**Y**) and *q*_*ψ,θ*_(**Z**^1:*P*^ |**X, Y**) denote the encoder posterior and the GP-updated posterior, respectively, and 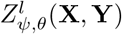 denotes the marginal likelihood of the *l*th latent dimension in the GP regression.

The key challenge in the above GP KL-divergence term is the computation of the marginal likelihood 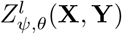 in the GP regression. According to [43], the evidence of the sparse GP regression for the *l*th latent dimension (in the first *P* dimensions) over all *N* data points is lower-bounded as:

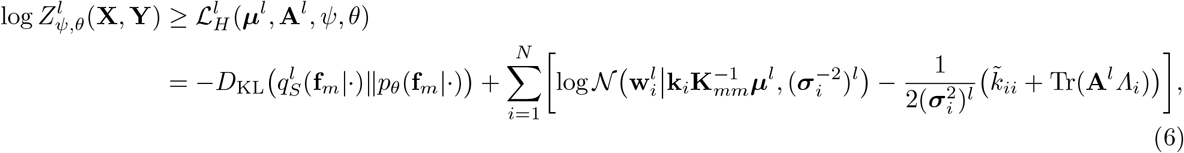

where 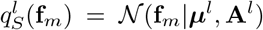 and *p*_*θ*_(**f**_*m*_) = 𝒩 (**f**_*m*_|0, **K**_*mm*_), **k**_*i*_ is the *i*th column of the matrix **K**_*mN*_, 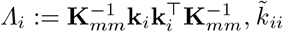 is the *i*th diagonal element of 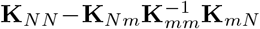 . Following [44,41], stochastic estimates of ***µ***^*l*^ and **A**^*l*^ for a single mini-batch *b* can be analytically computed as follows:

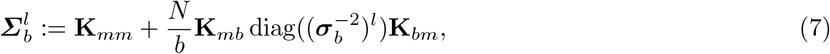

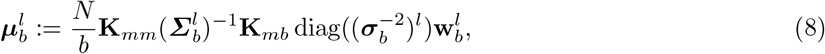

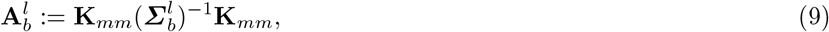

Then, the stochastic estimate of the posterior distribution of the latent embedding **z**^*l*^ based on mini-batch

*b* is:

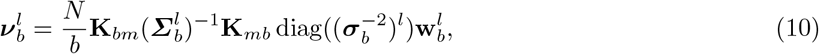

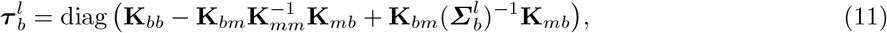

By combining all *P* latent dimensions, the posterior of the GP latent embedding based on mini-batch *b* is:

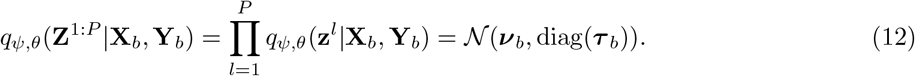

Finally, following [41], the mini-batch objective for the mini-batch *b* is:

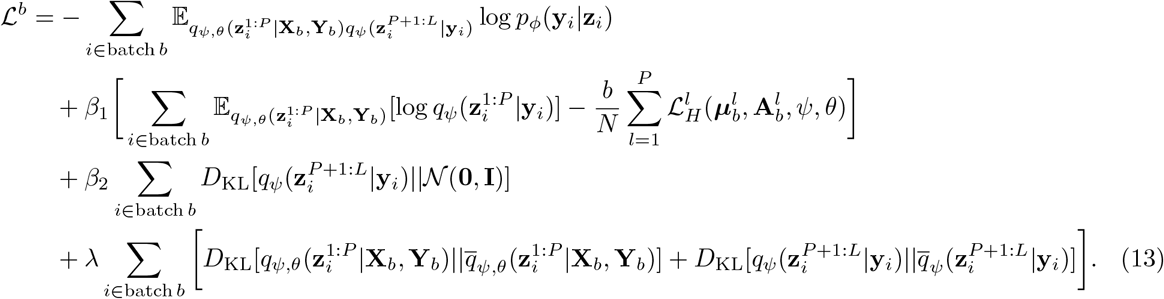

