## Supplementary notes and supplementary figures for "Decoupling Lineage and Intrinsic Information in Single-Cell Lineage Tracing Data with Deep Disentangled Representation Learning"

### Supplementary Notes 1: Datasets

#### 1 TedSim simulated data

We used TedSim [1] to simulate the cell division process from root cells, generating both gene expression profiles and lineage barcodes for each cell. The simulation constructs a binary lineage tree that records all division events. To reflect cellular heterogeneity, an asymmetric division mechanism [2,3,4] was incorporated, enabling one daughter cell to maintain the parental state while the other transitions to a new state. The evolution of cell states follows a predefined state transition tree and is regulated by the asymmetric division probability parameter $p_a$. Higher values of $p_a$ promote more frequent state transitions along the lineage. In this study, we evaluated four levels of $p_a$ (0.2, 0.4, 0.6, and 0.8) to analyze its influence on the resulting data.

The simulation employs a state identity vector (SIV) for each cell state, partially generated via a Brownian Motion Tree Model (BMTM) and supplemented with independent Gaussian noise. During division, each cell receives a cell identity vector (CIV) determined by:

$$\text{CIV}(\text{child}) = \text{CIV}(\text{parent}) + \text{SIV}(\text{child}) - \text{SIV}(\text{parent}) + \lambda\epsilon, \quad \epsilon \sim N(0, I).$$

Gene expression counts are then sampled following the SymSim framework [5], incorporating technical noise characteristic of single-cell sequencing protocols. We followed the TedSim to generate expression datasets of 500 genes for 4096 cells. This setup corresponds to a balanced binary lineage tree of depth 12 and was repeated for 4 different values of $p_a$. For lineage tracing, barcodes are modeled as CRISPR/Cas9-induced mutations, each consisting of 8 cassettes with 4 characters per cassette. Phylogenetic relationships are quantified using a scaled Hamming distance [6], which considers both the number of shared detectable sites and double weights transitions between "marked" and "unmarked" states.

#### 2 Mouse tumor dataset

The single-cell RNA sequencing data of mouse lung adenocarcinoma were obtained from the publicly available dataset published by [7], generated using the "KP-Tracer" model. This experimental system enables longitudinal tracing of tumor evolution from individual cells and provides phylogenetic trees, metadata, and gene expression matrices at single-cell resolution.

Two tumors were selected for analysis based on lineage scale and data completeness:

**3515_Lkb1_T1**: This KPL-type tumor exhibits a complex lineage structure and contains AT2-like cells along with a substantial proportion of pre-EMT cells. The sample includes matched primary and metastatic tumor data.

**3724_NT_T1**: Containing approximately 20,000 cells from multiple primary tumor regions and metastatic lesions, this sample displays pronounced cellular heterogeneity, with alveolar-like populations coexisting with cells transitioning toward invasive or mesenchymal-like states.

For data integration and preprocessing, lineage trees, metadata, and expression matrices were aligned using unique cell identifiers. Phylogenetic trees were pruned to retain only leaf nodes corresponding to cells present across all data types, with cells lacking expression data and single-node structures removed. All branch lengths were normalized to unity to ensure cross-sample comparability. Genes detected in fewer than 10 cells were filtered out, and the top 200 highly variable genes were selected for downstream analysis, reducing technical noise while preserving biologically relevant transcriptomic signals.

#### 3 Mouse ventral midbrain dataset

We selected the mouse ventral midbrain scRNA-seq dataset with endogenous lineage barcoding from [8], which comprises cells from embryonic day 11.5 (E11.5) and 15.5 (E15.5) for our integrated analysis.

The dataset captures a definitive developmental transition in cellular composition. At E11.5, the ventral midbrain was primarily composed of neural progenitors and neuroblasts, indicative of an active proliferative and fate-specification stage. By E15.5, the cellular landscape was dominated by differentiated neuronal subtypes, including dopaminergic (DA), glutamatergic (GLU), and GABAergic (GABA) neurons, marking a shift towards maturation.

A critical feature enabling our cross-temporal analysis is the stable and heritable nature of the CRISPR-generated mutational barcodes. The CREST system induces unique insertion/deletion (indel) patterns in the dual V1 and V2 recorder arrays within a short, early developmental window (by  E8.5). These barcodes are then permanently fixed and inherited, making the mutation information directly comparable between E11.5 and E15.5. An identical barcode in cells from both time points provides direct evidence of a shared clonal origin. By using the unique V1 and V2 barcodes to define clones, we screened the clones with a cell count of 6 or more, and finally obtained over eight thousand cells.

### Supplementary Notes 2: Evaluation Metrics

#### 1 Supervised Metrics

***Adjusted Rand Index (ARI)*** [9]: The ARI is used to quantify the agreement between a true partition $U = \{U_1, \dots, U_R\}$ and a clustering result $V = \{V_1, \dots, V_C\}$. Let $N$ be the total number of data points. An $R \times C$ contingency table $\mathbf{n}$ is constructed, where $n_{ij}$ denotes the number of elements shared between true class $U_i$ and cluster $V_j$. The marginal sums are $n_{i.} = \sum_j n_{ij}$ and $n_{.j} = \sum_i n_{ij}$. The ARI is defined as:

$$\text{ARI} = \frac{O - E}{M - E}$$

where the observed index ($O$), expected index ($E$), and maximum index ($M$) are calculated using the generalized binomial coefficient $\binom{x}{2} = \frac{x(x-1)}{2}$:

$$O = \sum_{i=1}^{R} \sum_{j=1}^{C} \binom{n_{ij}}{2}$$

$$E = \frac{\left(\sum_{i=1}^{R} \binom{n_{i.}}{2}\right) \left(\sum_{j=1}^{C} \binom{n_{.j}}{2}\right)}{\binom{N}{2}}$$

$$M = \frac{1}{2} \left[ \sum_{i=1}^{R} \binom{n_{i.}}{2} + \sum_{j=1}^{C} \binom{n_{.j}}{2} \right]$$

The terms $\sum_{i=1}^{R} \binom{n_{i.}}{2}$ and $\sum_{j=1}^{C} \binom{n_{.j}}{2}$ represent the number of pairs of elements placed in the same true class and the same cluster, respectively. ARI ranges from $-1$ to 1, with 1 indicating perfect concordance and 0 representing random agreement.

***Normalized Mutual Information (NMI)*** [10]: The NMI quantifies the mutual dependence between a true partition $U$ and a clustering result $V$. The calculation relies on the estimated joint probability $P(U_i, V_j)$ and marginal probabilities $P(U_i)$ and $P(V_j)$ from the contingency table:

$$P(U_i, V_j) = \frac{n_{ij}}{N}, \quad P(U_i) = \frac{n_{i.}}{N}, \quad P(V_j) = \frac{n_{.j}}{N}$$

The mutual information $I(U; V)$ is defined as:

$$I(U; V) = \sum_{i=1}^{R} \sum_{j=1}^{C} P(U_i, V_j) \log_2 \left( \frac{P(U_i, V_j)}{P(U_i) P(V_j)} \right)$$

The entropies $H(U)$ and $H(V)$ are:

$$H(U) = -\sum_{i=1}^{R} P(U_i) \log_2 P(U_i), \quad H(V) = -\sum_{j=1}^{C} P(V_j) \log_2 P(V_j)$$

The NMI, normalized by the average entropy (specifically the arithmetic mean normalization), is calculated as:

$$\text{NMI}_{\text{AM}}(U, V) = \frac{I(U; V)}{\frac{1}{2}[H(U) + H(V)]}$$

NMI ranges from 0 (no mutual information) to 1 (perfect correlation). Other normalization methods (e.g., geometric mean, max) also exist.

#### 2 Unsupervised Metrics

***Silhouette Coefficient*** [11]: The Silhouette Coefficient ($S$) measures how well each data point lies within its cluster. Let $C_I$ be the cluster containing data point $i$. The metric relies on two values for each point $i$:

$a(i)$: The average distance from $i$ to all other points in the same cluster $C_I$ (cohesion):

$$a(i) = \frac{1}{|C_I| - 1} \sum_{j \in C_I, i \neq j} d(i, j)$$

$b(i)$: The minimum average distance from $i$ to all points in any other cluster $C_K$ ($K \neq I$) (separation):

$$b(i) = \min_{K \neq I} \left\{ \frac{1}{|C_K|} \sum_{j \in C_K} d(i, j) \right\}$$

The Silhouette Coefficient for a single point $i$ is defined as:

$$s(i) = \frac{b(i) - a(i)}{\max\{a(i), b(i)\}}$$

The overall Silhouette Coefficient ($S$) for the clustering is the mean $s(i)$ over all $N$ data points:

$$S = \frac{1}{N} \sum_{i=1}^{N} s(i)$$

The coefficient ranges from $-1$ to 1. Values near 1 indicate dense and well-separated clusters; values near 0 suggest overlapping clusters; and negative values indicate that points might be assigned to the wrong cluster.

***Calinski-Harabasz Score*** [12]: The Calinski-Harabasz Score ($CH$) is defined as the ratio of the between-cluster dispersion mean and the within-cluster dispersion mean. Let $N$ be the total number of data points, $K$ be the number of clusters, $C_k$ be the set of points in cluster $k$, $n_k$ be the number of points in cluster $k$, $\mathbf{c}_k$ be the centroid of cluster $k$, and $\mathbf{c}_E$ be the overall centroid of the data.

$$CH = \frac{\text{Trace}(\mathbf{SS}_B)/(K-1)}{\text{Trace}(\mathbf{SS}_W)/(N-K)} = \frac{\text{SS}_B/(K-1)}{\text{SS}_W/(N-K)}$$

The between-cluster dispersion matrix ($\mathbf{SS}_B$) and the within-cluster dispersion matrix ($\mathbf{SS}_W$) are:

$$\mathbf{SS}_B = \sum_{k=1}^{K} n_k (\mathbf{c}_k - \mathbf{c}_E)(\mathbf{c}_k - \mathbf{c}_E)^T$$

$$\mathbf{SS}_W = \sum_{k=1}^{K} \sum_{\mathbf{x} \in C_k} (\mathbf{x} - \mathbf{c}_k)(\mathbf{x} - \mathbf{c}_k)^T$$

The scalar terms $\text{SS}_B = \text{Trace}(\mathbf{SS}_B)$ and $\text{SS}_W = \text{Trace}(\mathbf{SS}_W)$ represent the sum of squared distances for between-cluster and within-cluster dispersion, respectively, typically using the squared Euclidean distance. Higher $CH$ values indicate a better clustering result.

***Davies-Bouldin Index*** [13]: The Davies-Bouldin Index (DBI) is calculated based on the average similarity ($R_{ij}$) between each cluster $i$ and its most similar cluster $j$. Let $K$ be the number of clusters, $C_i$ be the set of points in cluster $i$, and $\mathbf{c}_i$ be the centroid of cluster $i$.

The measure of scatter within cluster $i$, $s_i$, is typically the square root of the average squared Euclidean distance (RMS error) to the centroid:

$$s_i = \left( \frac{1}{|C_i|} \sum_{\mathbf{x} \in C_i} \|\mathbf{x} - \mathbf{c}_i\|^2 \right)^{1/2}$$

where $\|\mathbf{x} - \mathbf{c}_i\|$ is the Euclidean distance between point $\mathbf{x}$ and cluster centroid $\mathbf{c}_i$.

The distance between the centroids of cluster $i$ and cluster $j$ is denoted by $d_{ij}$ (usually Euclidean distance):

$$d_{ij} = \|\mathbf{c}_i - \mathbf{c}_j\|$$

The similarity (or cluster separation ratio) between cluster $i$ and cluster $j$ is defined as:

$$R_{ij} = \frac{s_i + s_j}{d_{ij}}$$

For each cluster $i$, the highest similarity to any other cluster is found:

$$D_i = \max_{j \neq i} R_{ij}$$

The overall Davies-Bouldin Index (DBI) is the average of these maximum similarities across all clusters:

$$\text{DBI} = \frac{1}{K} \sum_{i=1}^{K} D_i$$

Lower values of DBI indicate better clustering, signifying good separation between clusters and high density within clusters.

### Supplementary Figures

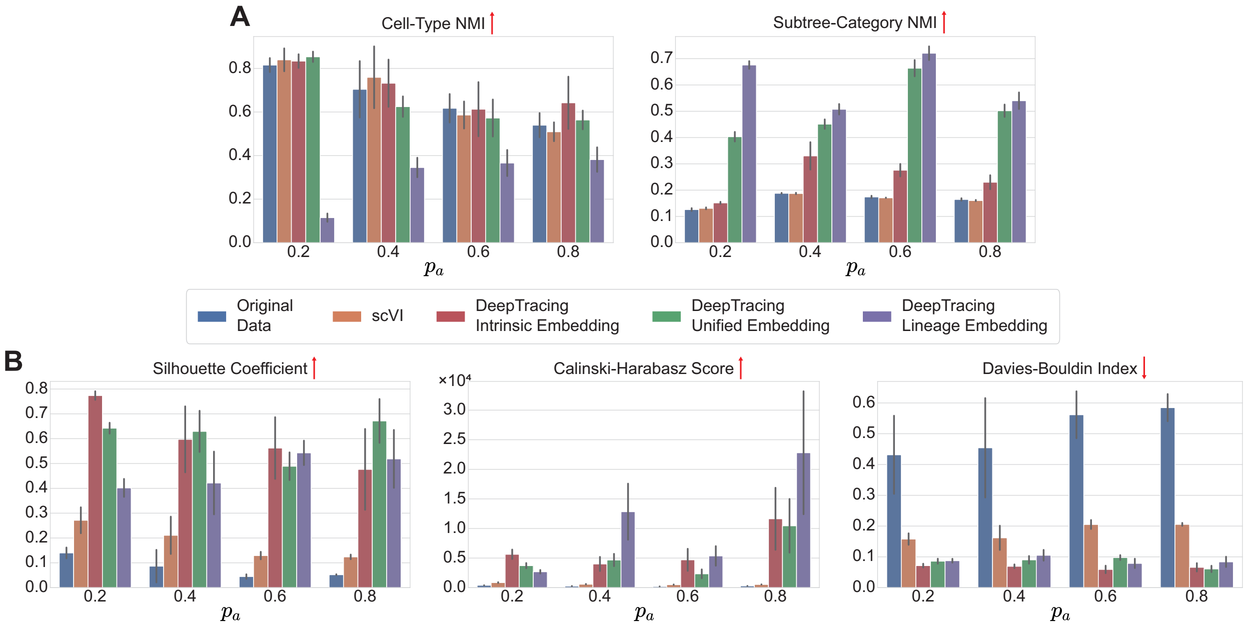


**Supplementary Figure 1:** Valuation of different embeddings across various values of the simulation parameter $p_a$. (A) Supervised metrics: cell-yype NMI and subtree-category NMI. (B) Unsupervised metrics: Silhouette coefficient, Calinski-Harabasz score, and Davies-Bouldin index.

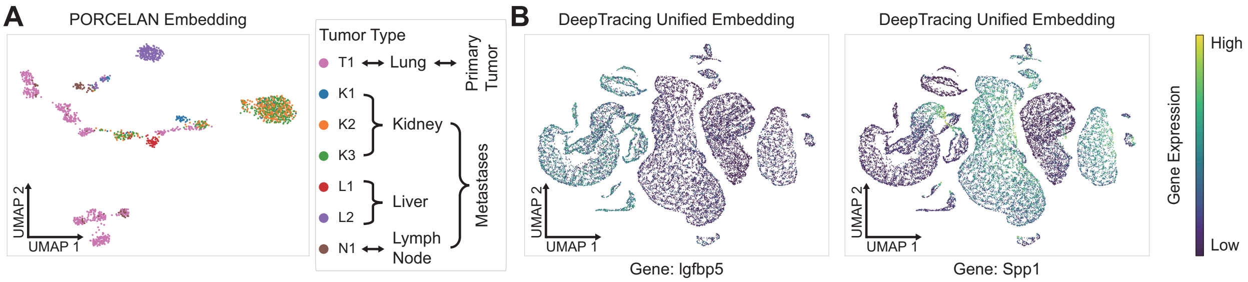


**Supplementary Figure 2:** (A) UMAP visualization of the PORCELAN embedding, colored by tumor type. (B) UMAP visualization of the DeepTracing unified embedding, colored by expression levels of two key genes: *Igfbp5* (left) and *Spp1* (right).
